# Transcriptional analysis supports the expression of human snRNA variants and reveals U2 snRNA homeostasis by an abundant U2 variant

**DOI:** 10.1101/2020.01.24.917260

**Authors:** Brian Kosmyna, Varun Gupta, Charles Query

## Abstract

Although expansion of snRNA genes in the human genome and sequence variation in expressed transcripts were both identified long ago, no study has comprehensively analyzed which genes are transcriptionally active. Here, we use comprehensive bioinformatic analysis to differentiate between similar or identical genomic loci to determine that 49 snRNA genes are actively transcribed. This greatly expands on previous observation of sequence variation within snRNA transcripts. Further analysis of U2 snRNA variants reveals sequence variation maintains conserved secondary structures, yet sensitizes these U2 snRNAs to modulation of assembly factors. Homeostasis of total U2 snRNA level is maintained by altering the ratio of canonical and an abundant U2 snRNA variant. Both canonical and variant snRNA promoters respond to MYC and appear differentially sensitive to increased MYC levels. Thus, we identify transcribed snRNA variants and the sequence variation within, and propose mechanisms of transcriptional and post-transcriptional regulation of snRNA levels and pre-mRNA splicing.

**HIGHLIGHTS:**

- ChIP-seq of active promoters identifies uncharacterized snRNA genes
- Transcribed repetitive snRNA genes are distinguished from falsely-mapped snRNA loci
- U2 snRNA variants are sensitive to modulations in snRNP assembly
- Widely expressed U2 snRNA variants provide homeostasis for total U2 snRNP levels

## INTRODUCTION

Intron removal from nascent pre-mRNA is an essential step of RNA processing and gene regulation in eukaryotes. The two catalytic steps of splicing, 5’ splice site (SS) cleavage/lariat branching and 3’ SS cleavage/exon ligation, are catalyzed by the spliceosome, a large, dynamic complex that comprises small nuclear ribonucleoprotein complexes (snRNPs), which themselves contain five small nuclto ensure precise excision of each intron (reviewed in Will and Luhrmann, 2011). Throughout these transitions between spliceosomal complexes, snRNAs make many critical contacts either through RNA-RNA or RNA-protein interactions. The importance of these interactions can be inferred from evolutionary conservation of sequence motifs and secondary structures formed by snRNAs and have been confirmed through biochemical and genetic experiments (reviewed in Mayerle and Guthrie, 2017; Will and Luhrmann, 2011).

Recent cryo-EM structures of the spliceosome have provided both atomic detail and overall molecular architecture of many spliceosomal complexes (Kastner et al., 2019; Plaschka et al., 2019; Yan et al., 2019). These structures reveal massive architectural dynamics of U2 snRNP, whereas U5 and U6 snRNPs remain relatively static through spliceosomal activation, catalysis, and disassembly (Kastner et al., 2019; Plaschka et al., 2019; Yan et al., 2019). For example, when the U2 snRNA GUAGUA motif base pairs to the intron branch site early in spliceosome assembly, it forms the U2:branchsite (U2:BS) duplex, which is protected by the HEAT domains of SF3B1 (Plaschka et al., 2018). This interaction sequesters the hydroxyl group of the bulged adenosine, the first-step nucleophile, from the 5’SS and prevents formation of an active catalytic core. Release of SF3B1 from the Bact complex allows U2 snRNA to undergo a large rotation that collapses the U2:BS duplex into the active site and positions the reactants for first-step catalysis (Galej et al., 2016; Haselbach et al., 2018; Yan et al., 2016; Zhan et al., 2018). In addition, U2 snRNA forms dynamic secondary structures with itself and with U6 snRNA that are essential to multiple steps of the splicing cycle (Hilliker et al., 2007; Mefford and Staley, 2009; Perriman and Ares, 2010; Perriman and Ares, 2007).

Whereas functional roles of each snRNA have been elucidated through studies in human and yeast, recently it has been suggested that snRNA expression levels themselves may regulate alternative splicing in human tissues and correlate with changes observed in disease (Dvinge et al., 2019). Early work to identify and characterize snRNA genes revealed that U1 and U2 snRNA genes are present as both large multi-copy clusters and pseudogenes of similar sequence spread throughout the genomes of many model organisms (Denison et al., 1981; Denison and Weiner, 1982; Lindgren et al., 1985; Lund and Dahlberg, 1984; Manser and Gesteland, 1982; Van Arsdell and Weiner, 1984). Later, conserved sequence elements were discovered in the promoters of all canonical snRNA genes, and all major snRNAs were discovered to be transcribed by RNA polymerase II (Pol II), except U6 snRNA which is transcribed by RNA polymerase III (reviewed in Didychuk et al., 2018; Guiro and Murphy, 2017). Identification of sequence heterogeneity of U1, U4, and U5 suggested that snRNA variants could be differentially expressed, and it was hypothesized that expression of low abundance snRNAs may regulate alternative splicing (Branlant et al., 1983; Forbes et al., 1984; Krol et al., 1981; Lund, 1988; Lund and Dahlberg, 1987; Sontheimer and Steitz, 1992). Therefore, identifying low abundance snRNAs that affect alternative splicing could provide insight into how endogenous nucleotide variation effects splicing mechanisms.

In addition to sequence heterogeneity found in many metazoans, a second spliceosome, the minor or U12 spliceosome, excises a limited number of introns with non-canonical splice site sequences. The minor spliceosomal U11, U12, U4atac, and U6atac snRNAs are analogous to snRNAs in the major U2 spliceosome, and U5 snRNA is common to both (Patel and Steitz, 2003; Turunen et al., 2013). Although the snRNA sequences are greatly divergent, the minor spliceosome could be considered an example of low abundance snRNA variants affecting the splicing of specific introns. Recent knock down of a variant U1, RNVU1-8 expressed in HeLa cells, altered levels of specific transcripts, whereas its ectopic expression in human skin fibroblasts led to an increase in two critical stem cell markers, Nanog and SOX2 (O’Reilly et al., 2013; Vazquez-Arango et al., 2016). Despite being proposed at the time of the discovery of snRNA variants, such investigation into mechanisms through which variant snRNAs might regulate alternative splicing has been limited.

Although technical advances have been made in sequencing and detection of rare sequence variation in both DNA and RNA transcripts, a complete understanding of snRNA variant expression has yet to be achieved. Many RNA-seq protocols exclude abundant small RNAs before library preparation due to length. Further, if reads that align to snRNAs are present in RNA-seq libraries, they are bioinformatically excluded because their sequences match many loci in reference genomes, or they are assigned randomly to one of these many loci. These biases of library and bioinformatic exclusion prevent accurate detection and quantification of both canonical snRNAs and their variants. Further complicating the analysis of snRNAs in previous sequencing experiments is the copy number variation of the canonical U2 snRNA, which was excluded from the hg19/GRC37 human reference genome, to which the majority of extant RNA-seq data has been aligned (Tessereau et al., 2013; Tessereau et al., 2014; Westin et al., 1984).

Here, we present analyses of ENCODE ChIP-seq experiments that provide evidence for the expression of novel snRNA variants, including expression of twenty-four U1 and four U2 snRNA genes with unique mature sequences. We further investigated potential functions of novel U2 snRNA variants. We detected the expression of one major U2 snRNA variant that is abundant in human tissues and two minor variants that are less abundant. All secondary structures are maintained in these variants with either compensatory or conservative Watson-Crick (W-C) or wobble base-pair changes. Although canonical U2 snRNA (RNU2-1, hereafter U2-1) and the next-most abundant U2 snRNA, RNU2-2 (hereafter U2-2) are expressed at widely different ratios in a tissue-specific manner, MCF7 U2-2^-/-^ cells generated with CRISPR/Cas9 are viable with no significant phenotype. The upregulation of U2-1 snRNA in the absence of U2-2 maintains pre-mRNA splicing and gene expression in these cells and suggests that homeostasis of U2 snRNA levels is provided by variant U2-2 expression. Rescue of U2-2 expression by both canonical U2-1 and variant U2-2 promoters in MCF7 U2-2^-/-^ cells further supports that regulation of total U2 snRNA occurs post-transcriptionally. Overall, this report provides strong evidence for expression of variants of all major snRNAs, post-transcriptional regulation of U2 snRNA, and homeostasis of total U2 snRNA level by expression of U2 variants.

## RESULTS

### Expansion of snRNA “pseudogenes” in higher eukaryotes

Genomic sequencing of many eukaryotic model organisms confirmed early observations that snRNA sequences are highly repetitive. However, comparisons between species reveals a correlation between genome complexity (i.e. size, number of genes, and the prevalence of alternative splicing) and the number of annotated snRNA genes and pseudogenes (**Table S1**) (Kalvari et al., 2018; Zerbino et al., 2018). In *S. cerevisiae*, there are a total of five snRNA genes (one copy of each snRNA in the major spliceosome), whereas *D. melanogaster* and *C. elegans* have 47 and 79 annotated snRNA genes, respectively. This expansion of snRNA genes is more drastic in vertebrates: *D. rerio*, *M. musculus*, and *H. sapiens* have 1160, 1296, and 1865 annotated snRNA genes and pseudogenes, respectively (**Figure 1A**).

**Figure 1.**
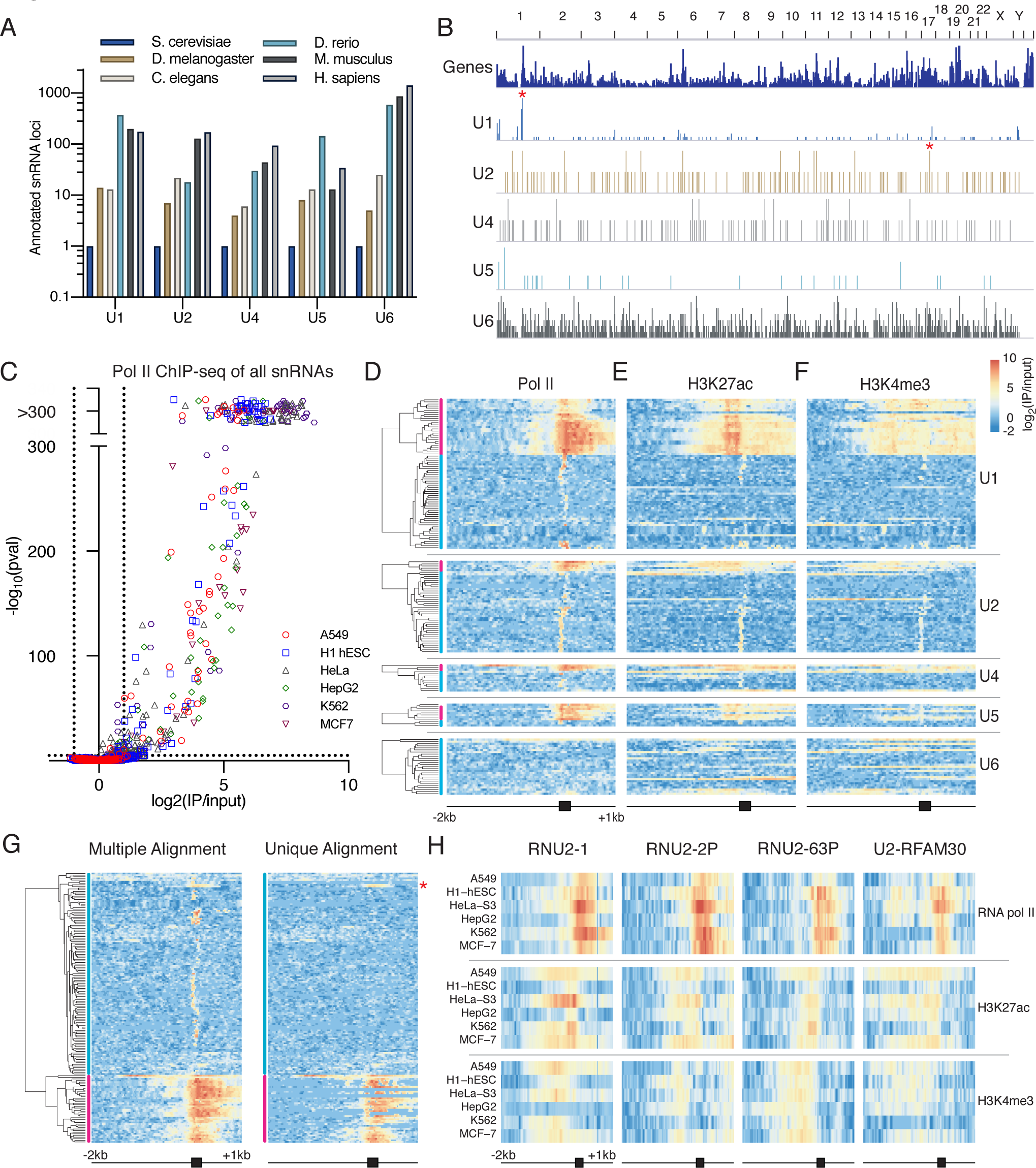
Prevalence, distribution, and RNA Pol II ChIP of annotated snRNA genes. (A) Prevalence across model organism genomes. Each bar represents the number of snRNA genes or pseudogenes annotated by Gencode or Rfam (Kalvari et al., 2018; Zerbino et al., 2018) for each of the snRNAs in the major spliceosome for select eukaryotic model organisms. Scale is in log10. (B) Genomic distribution of all annotated human snRNA genes and pseudogenes. IGV used to visualize tracks containing snRNA genes are shown. Chromosome number is listed along top of genome browser; each vertical line represents the presence of snRNA genes or pseudogenes within a minimally resolvable distance. The y-axis scale varies between each snRNA and is determined by the number of snRNA genes in each bin. The multi-copy arrays of canonical U2-1 on chr17 and U1-1 on chr1 are indicated by red (*). The canonical U2-1 array is shown as a single gene. (C) Volcano plot of significant enrichment of Pol II density over each snRNA gene in six cell lines. log2(IP/input) on the x-axis shows magnitude of enrichment while -log10(p-val) on the y-axis is calculated from a binomial test. All -log10(p-val) greater than 300 are shown above the break in graph; jitter (noise) was added to these values for visualization. Cut-offs for significance are indicated by dotted lines. (D) Heatmaps of ENCODE ChIP-seq data in MCF7 cells. Reads were mapped to hg38 and allowed to align to up to 20 positions. The U2-1 cluster was masked, and a single gene repeat was added to hg38 as a contig. Each row represents a different snRNA gene from distinct genomic loci. Dendogram from hierarchical clustering of each snRNA and designation of class I/actively transcribed (pink bar) and class II/inactive (blue bar) snRNA genes are shown on left. All snRNAs in heatmaps are significantly enriched for Pol II density in at least one of the six cell lines analyzed. Each row includes 2 kb upstream and 1 kb downstream relative to the TSS. The TSS and approximate length of a mature snRNA (200 bp) are indicated by the black box. (E, F) Heatmap of ENCODE ChIP-seq data of H3K27ac and H3K4me3 in MCF7 cells. Each row represents a different snRNA gene from distinct genomic loci and snRNA genes are in the same order as in (D). Scale shown is for all heatmaps in Figure 1. (G) Comparison of snRNA genes with multiple alignment (>20 loci) or unique alignment which discards reads aligning to more than one genomic position. All 167 snRNAs from (D-F) shown. Hierarchical clustering was performed on multiple alignment (left) and order of rows was maintained for unique alignment (right). * indicates cluster of class I genes from (D). (H) Multiple alignment of ChIP-seq Pol II, H3K27ac, and H3K4me3 of transcribed U2 snRNA genes. See also **Figure S1**.

In the human genome, snRNA genes are present on every chromosome as multicopy arrays or single genes (**Figure 1B**, **Table S2**). The organization and sequence characteristics of snRNA genes and pseudogenes support an early proposal that these copies were generated through distinct mechanisms. The resulting genes could have varied genomic environments, some with promoter and downstream sequences similar to the canonical genes, others with similar mature snRNA sequences but diverse promoters and downstream sequences, and lastly some pseudogenes with high sequence similarity to the 5’ ends of the mature snRNA but no further sequence similarity (Denison and Weiner, 1982). Although expansion of snRNA genes in the genome and sequence variation in expressed transcripts were both identified long ago, no study has comprehensively analyzed which genes are transcriptionally active.

### Detecting transcription of repetitive snRNA genes

Although the detection of sequence variation in the genome and transcriptome has improved with sequencing technologies and bioinformatic analyses, many RNA-seq protocols exclude small abundant RNA species while enriching for polyadenylated mRNAs. Furthermore, fragments aligning to repetitive sequences in a reference genome are often bioinformatically discarded. The probability of a sequence fragment aligning uniquely to a repetitive genomic element increases with longer fragment length and higher sequence variability within a repetitive element. However, the canonical sequences for human U1 and U2 snRNAs are present as near-identical copies in arrays on chromosomes 1 and 17, respectively (asterisks in **Figure 1B**). While some bioinformatic approaches exclude these reads from further analysis, others misalign them to identical regions of snRNA genes that have no other evidence they are actively transcribed. Combined, these challenges have prevented accurate detection and quantification of expressed snRNA genes.

To determine which snRNA variants are transcribed, we analyzed ChIP-seq of Pol II and two histone modifications indicative of active transcription, H3K27ac and H3K4me3, from ENCODE (Davis et al., 2018). In contrast to RNA-seq data, ChIP-seq data allows for alignment to the divergent sequences of promoters and regions downstream from the highly similar mature snRNAs. This increases the probability of uniquely identifying the genomic origin of each fragment. Additionally, comparisons of multiple datasets in multiple cell lines increase confidence that specific snRNA genes are transcribed. We first aligned reads from ENCODE ChIP-seq experiments for Pol II, H3K27ac and H3K4me3 in six common cell lines to the human genome allowing reads to align to a maximum of 20 loci with no mismatches. As a control and to assign areas to determine enrichment, canonical U1 and U2 snRNA genes were analyzed for regions of positive ChiP-seq signal (H3K27ac from -400 bp to +200 bp, H3K4me3 from -1000 bp to +200 bp, and Pol II from -50 bp to +400 bp relative to the transcription start site (TSS)) (**Figure S1A)**. These regions of positive signal on the canonical U1 and U2 genes were then used to calculate ChIP-seq signal (log2(IP/input)) and enrichment score (-log10(p-value)) for all annotated snRNA genes. Enrichment of all snRNA genes in each cell line is shown for Pol II, H3K27ac, and H3K4me3 (**Figures 1C**, **S1B**, and **S1C**). 167 snRNA genes displayed significant enrichment of Pol II density in at least one of the six analyzed cell lines.

Hierarchical clustering of these 167 snRNA genes by Pol II ChIP-seq signal revealed two major classes of enriched snRNA genes (**Figure 1D**). This clustering differentiates between expressed snRNA genes with strong signal and enrichment across the entire region observed in canonical genes from those that only have enrichment over a fraction of the region interrogated for significant ChIP-seq signal. While transcription of most loci in the first class is supported by enrichment of both H3K27ac and H3K4me3 (**Figure 1E**, **1F**, and **Table S3**), histone modifications indicative of active promoters, the second class lacks enrichment of these modifications suggesting that apparent enrichment on the genes making up the second class could be an artifact of multiple alignment. To test this, we compared the multiple alignment of all 167 Pol II-enriched snRNA genes to an alignment that only mapped unique reads with zero mismatches (**Figure 1G** and **Table S3**). For most genes in the second class, all aligned reads exactly match another expressed gene, suggesting they originate from a different locus. However, a small fraction of class II snRNA genes were still significantly enriched for Pol II density when only uniquely aligned reads were considered. Since these unique reads are restricted to the mature but not flanking regions of snRNA genes and there is often no enrichment of H3K27ac and H3K4me3, it is likely these genes are not expressed. These aligned reads may be due to sequencing error, presence of a single nucleotide polymorphism (SNP), or bioinformatic misalignment due to trimming (“soft clipping”) or insertions/deletions to improve the read alignment. This analysis provides strong evidence of transcription for some snRNA genes (class I), while also determining which snRNA genes are prone to false identification due to their repetitive sequence and bioinformatic misalignment (class II).

We found strong evidence for transcription of thirty-two U1, four U2, two U4, and eight U5 snRNA genes including all the canonical snRNA genes for the major and minor spliceosome. U6 snRNAs were not identified since they are not transcribed by Pol II. Eight of the transcribed U1 genes have mature sequences identical to the canonical U1 snRNA gene, but variation in the flanking sequences and mapping reads to multiple genomic loci allowed some to be identified as transcribed with high confidence. We identified transcribed genes with sequence variation that match 6 of 8 previously identified variant U1 transcripts (O’Reilly et al., 2013; Somarelli et al., 2014). We also identified two transcribed genes from the variant U1 cluster that were previously examined and thought not to be transcribed. Taken together, our genome-wide analysis in multiple cell lines differentiates between highly similar or identical genomic loci and identifies those with active promoters.

Our analysis suggests four U2, two U4, and eight U5 snRNA genes are transcribed in multiple cell lines (**Figure 1H**, **S1D**, and **S1E**). Sequence variation of both U4 and U5 snRNAs has previously been observed in snRNA transcripts (Bark et al., 1986; Sontheimer and Steitz, 1992). However, two of the U2 snRNA variants have never been reported as transcribed, and the U2-2 snRNA has often been reported bioinformatically as “U2” due to its similarity in sequence to U2-1 and to U2-1’s absence from previous genome assemblies (Nojima et al., 2018; e.g., Nottingham et al., 2016). Since sequence variation in U2 snRNA has not been previously described in humans and considering the multiple roles U2 snRNA has in splice site selection, formation of the catalytic core, and presentation of the pre-mRNA substrates into the active site, we further investigated the transcribed U2 snRNA variants.

### Differential variant U2 snRNA expression

Variant-specific primers were designed for numerous U2 snRNAs and their specificity was confirmed by their inability to cross amplify other variants, either from bacterial artificial chromosomes (BACs) containing only a single variant or from a synthetic double-stranded DNA template (**Figure S2A**). The identities of qPCR amplicons from RNA samples were confirmed by Sanger sequencing. We then determined the expression levels of three of four U2 snRNAs enriched for RNA pol II density, H3K27ac, and H3K4me3 from the previous analysis in diverse tissue types. The canonical U2-1 and similar variant U2-2 snRNA are highly abundant and represent the majority of total U2 snRNA in all RNA samples tested from different human tissues (**Figure 2A**). Therefore, we compared the levels of other variant U2 snRNAs, including minor spliceosomal U12, to the combined level of U2-1 and U2-2. The ratio of U2-2 to total U2 in all tissues except fetal heart ranges from approximately 12% to 43%, whereas in fetal heart U2-2 represents the majority of total U2 snRNA (**Figure 2A**). In these RNA samples from human tissues, U12 also varies significantly with all but two samples between 4.4% and 17%, whereas breast tumor and testis represent outliers of the distribution with 28% and 38%, respectively. Although U12 levels have historically been thought to be near 1% of U2 snRNA, our data suggest U12 snRNA expression can represent a larger fraction when compared to U2 snRNA and the ratio of U12 to U2 can vary in different tissues. Of the two more divergent U2 snRNAs, U2-63 was detected in the majority of tissues at between ∼1% and 0.1% of total U2 snRNA (**Figure 2B**), whereas no primers efficiently amplified sequences for U2-R30. This ratio, although low, may represent a significant number of transcripts due to the high total expression of U2 snRNA (approximately 500,000 transcripts per cell (Baserga and Steitz, 1993)).

**Figure 2.**
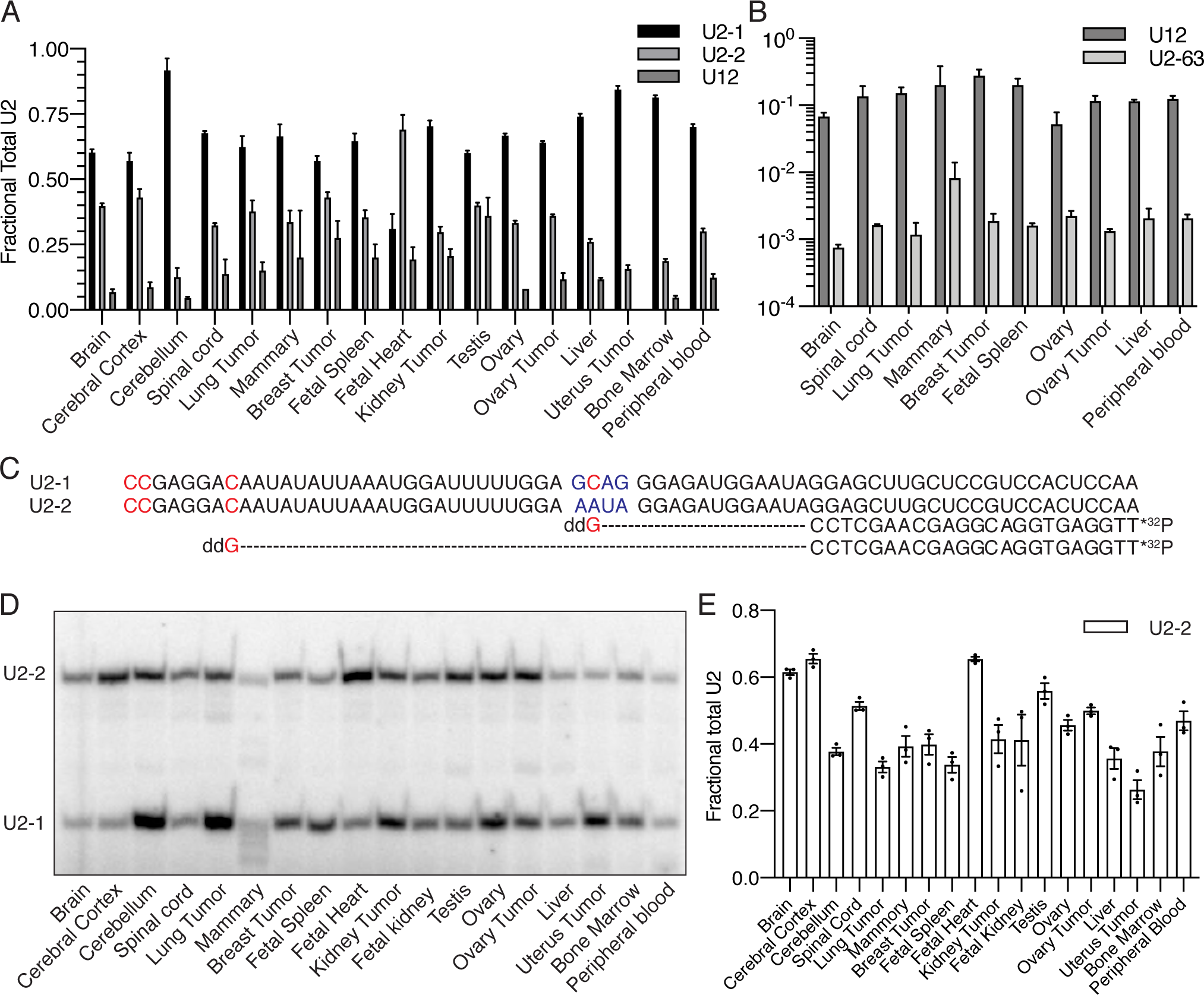
Expression of human U2 snRNAs. (A) Relative abundance of U2-1, U2-2, and U12 snRNAs in human tissues as detected by RT-qPCR using variant-specific RT-qPCR primers. Error bars represent SEM of three technical replicates human RNA samples. (B) Relative amount of U12 and U2-63 snRNAs in same human tissue samples as detected by RT-qPCR using variant-specific RT-qPCR primers. Error bars represent SEM of three technical replicates of human RNA samples. Scale is log10. (C) Experimental design of variant-specific detection of U2-1 and U2-2 by primer extension with products from U2-1 and U2-2 snRNAs shown below. (D) Primer extension confirms abundance of U2-2 in RNA from human tissue samples. (E) Quantification of percent U2-2 from primer extension as calculated by U2-2 intensity divided by sum of U2-2 and U2-1 intensity. Error bars represent SEM of three technical replicates of human RNA samples. See also **Figure S2**.

Since U2-1 and U2-2 represent the majority of U2 snRNA in human tissues and are highly similar, we developed a variant-specific primer extension assay in which a DNA oligonucleotide primer hybridized equally well to both variants. U2-1 and U2-2 were then differentiated by incorporation of ddGTP and termination of reverse transcription at different positions in the cDNA (**Figure 2C**). The cDNAs could then be separated and quantified to determine the ratio of U2-2 to total U2 snRNA levels (**Figure 2D)**. The primer extension assay confirms the ratio of variant U2-2 to canonical U2-1. However, primer extension detects higher U2-2 levels than RT-qPCR in 14 of 18 samples, and U2-2 is more abundant than U2-1 in 6 of 18 tissues tested (**Figures 2A** and **2E**). Although there are observed differences between the two assays, there is a significant correlation (**Figure S2B**) between primer extension and RT-qPCR suggesting the ratio of U2-1 and U2-2 is differentially regulated across diverse human tissues.

### Transcribed U2 snRNA variants maintain secondary structures

The U2 snRNA genes enriched for RNA pol II, H3K27ac, and H3K4me3 include the canonical U2-1, the highly similar variant U2-2, and two variants with many nucleotide differences spread throughout the mature sequence, U2-63 and U2-R30. U2-1 and U2-2 snRNA genes vary by five nucleotides, four consecutive nucleotides downstream of the Sm site before stem loop III and a single nucleotide difference in the loop of stem loop IV (**Figures 3A-3C** and **S3**). The sequences near the Sm site make no known specific contacts in the spliceosome, but are in the minimal motif needed for Sm ring assembly (Golembe et al., 2005). The two more divergent U2 snRNA variants, U2-63 and U2-R30, have 42 and 55 nucleotide differences when aligned and compared to U2-1 snRNA. However, most of these changes represent conservative or compensatory nucleotide substitutions that maintain secondary structure (**Figures 3C-3F**). Some of these secondary structures are formed transiently in specific spliceosomal complexes or must be altered during transitions between complexes, such as U2 stems IIa and IIc toggling between the two catalytic steps, and the opening of stem loop I for interaction with U6 snRNA during formation of Bact. Whereas these sequence variations may alter some interactions in or transitions between spliceosomal complexes, many interactions seem likely to be maintained in all four U2 snRNA variants, e.g. the highly conserved stem-loop IV that binds U2 snRNP components U2A’ and U2B” suggesting these interactions are unchanged.

**Figure 3.**
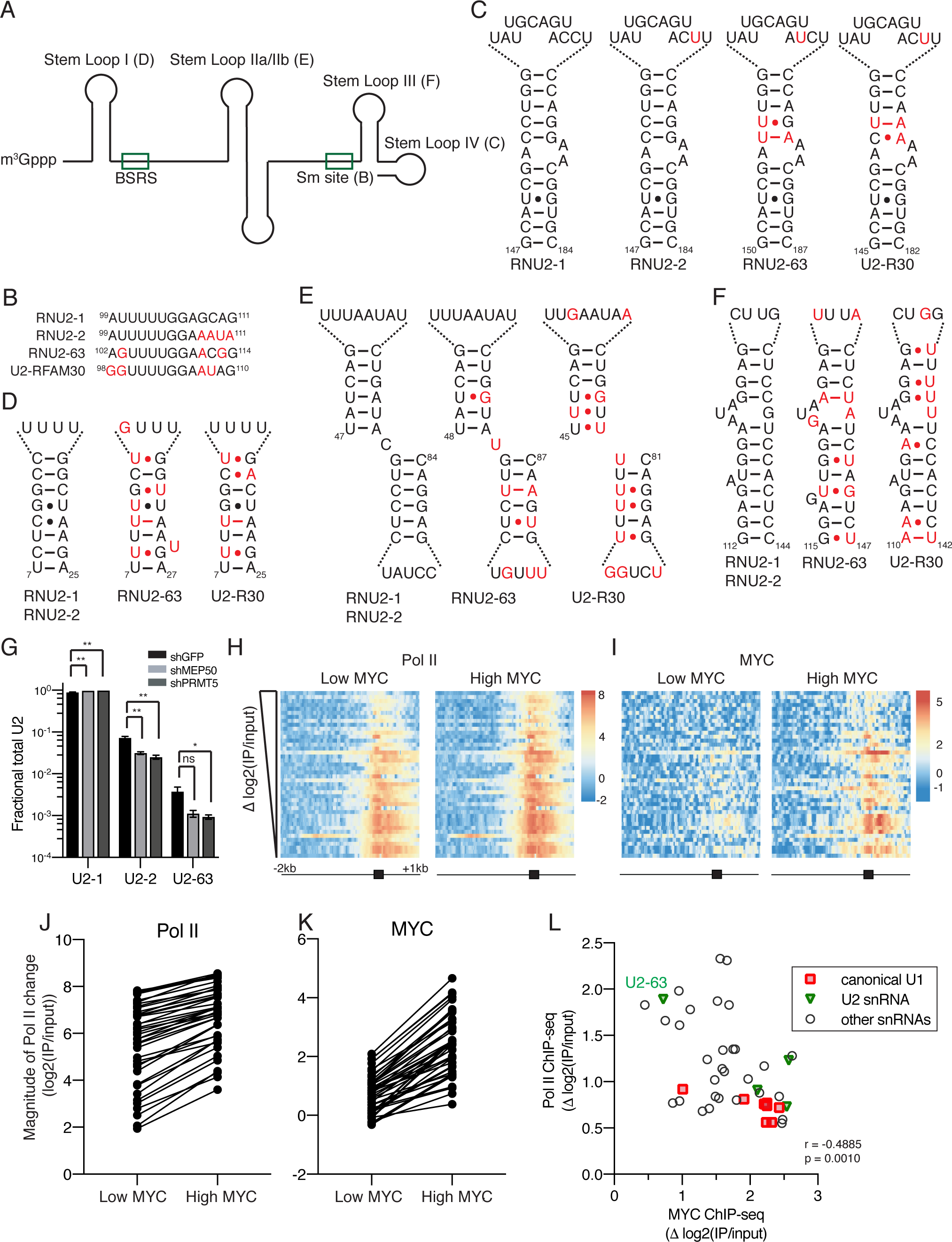
Modulation of snRNP assembly affects U2 variant levels. (A) Secondary structure of human U2 snRNA. Branchsite recognition sequence (BSRS) and Sm binding site (Sm Site) are indicated by green boxes. (B) Nucleotide variation between four expressed U2 variants maintains secondary structure in Sm site, nucleotide variation between U2 snRNAs is highlighted in red, basepairing interactions for Watson Crick interactions (–) or wobble interactions (•) are shown for (C-F). (C) Stem loop IV. (D) Stem loop I. (E) Stem loop IIa/IIb. (F) Stem loop III. (G) U2 snRNA levels in A549-shGFP, -shMEP50 and -shPRMT5 cell lines. Fractional Total U2 is displayed in log10 scale. Error bars represent SEM of biological replicates from three consecutive passages of the cell lines. P-value for U2-63 levels are 0.053 and 0.040, respectively. (H) ChIP-seq analysis of class I/transcribed snRNA genes in P324 cells upon endogenous MYC expression (low MYC) or overexpression (high MYC). Heatmap as in Figure 1 except ordered from greatest Pol II density increase upon MYC overexpression to least. (I) ChIP-seq of MYC binding to snRNA genes upon MYC overexpression. snRNA genes are in same order as in (H). (J-K) ChIP-seq signal (log2(IP/Input)) of Pol II and MYC binding upon MYC overexpression for each transcribed snRNA. (L) Correlation between magnitude of change in ChIP-seq signal (log2(IP/Input)) for Pol II and MYC upon MYC overexpression. r and p values were calculated using Pearson correlation. See also **Figure S3**.

### U2 variants are sensitive to modulation in snRNP assembly

Because U2 variants differ in sequence within and/or adjacent to the Sm binding site (**Figure 3B**), we hypothesized that they might be assembled with differing efficiencies. To test this, we used cells expressing shRNAs targeting two assembly factors, PRMT5 and MEP50, compared to shGFP as a control. We chose these two factors because they only mildly decrease assembly, allowing for cell viability, unlike other components of the assembly machinery, like SMN (Chen et al., 2017; Zhang et al., 2008). Steady-state U2 snRNA levels revealed a reduction in U2-2 and U2-63 compared to U2-1 in both knock-down cell lines relative to the shGFP control cell line (**Figure 3G**). Since snRNAs that are not properly assembled are thought to be quickly degraded (Sauterer et al., 1988; Shukla and Parker, 2014; Zieve et al., 1988), changes in the concentration of assembly, quality control, or degradation components could alter the pool of snRNAs available to be assembled and stabilized. We conclude that U2 variants are assembled less efficiently than U2-1, and when assembly components are limiting they are not stably incorporated into snRNPs. Thus, snRNP assembly, in particular Sm ring deposition, may be a significant control point for regulation of snRNA variant levels.

### MYC overexpression increases variant snRNA transcription

MYC overexpression in B-cell lymphomas upregulates snRNP assembly factors (Koh et al., 2015); we asked if this also results in upregulation of variant snRNAs. Using our bioinformatic pipeline to reanalyze ChIP-seq of Pol II and MYC from P493 cells with either low or high MYC expression (Sabo et al., 2014), we observed increased Pol II and MYC coverage on expressed snRNA genes when MYC is overexpressed (**Figure 3H**, **3I** and **Table S3**). snRNA genes with low Pol II density when MYC is not overexpressed had larger increases in Pol II density upon MYC overexpression, compared to snRNA genes with high Pol II density (**Figure 3H** and **3J**). Conversely, snRNA genes with low MYC occupancy in the low MYC condition displayed only slight increases in MYC binding upon overexpression, whereas genes displaying higher MYC binding in the low MYC condition showed the greatest increase in MYC density (**Figure 3I** and **3K**). Further, there is an inverse correlation between the amplitudes of change in ChIP signal in Pol II and MYC on snRNA genes **(Figure 3L)**. Genes with high Pol II density under low MYC conditions bind the most MYC as its concentration is increased, but this increase in MYC signal does not result in increased Pol II density. Conversely, genes with low Pol II density in low MYC conditions bind only slightly more MYC when it is upregulated, but this slight increase in MYC binding corresponds to large increases in Pol II density. While transcription of canonical U1 snRNA genes appears to only slightly increase upon MYC overexpression, transcription of some variant snRNAs including U2-63 appears more responsive to increases in MYC expression **(Figure 3L)**.

### Homeostasis of U2 snRNA levels maintained upon U2-2 knockout

Since U2-2 is highly expressed in most tissues, we utilized CRISPR/Cas9 to excise the mature snRNA sequence from the genome (**Figure S4A**). MCF7 cells were transfected with two plasmids, each containing Cas9, GFP, and a single gRNA targeting either upstream or downstream of U2-2. The transfected cells were GFP+ single-cell sorted and viable clones were expanded and genotyped. Clonal wild-type, heterozygous, and knock-out cell lines were isolated (**Figure 4A**). However, the presence of U2-2 was not detected by RT-PCR in heterozygous MCF7 clones with both WT and KO bands (**Figure 4B**). Sequencing of the wild-type sized band from heterozygous lines revealed a gene inversion for U2-2 mature sequence in all three samples shown. Primer extension was performed on the MCF7 U2-2^+/+^ parental cell line in addition to U2-2^+/+^, U2-2^-/-^ and U2-2^+/-^ clonal cell lines. While no differences in the ratio of U2-2 to U2-1 in the wild-type MCF7 cell lines were observed, cells lacking U2-2 show no U2-2 expression, confirming the specificity of both primer extension and RT-qPCR assays to differentiate between these two highly similar U2 snRNAs (**Figures 4C** and **S4B**). Interestingly, by primer extension we observed an up-regulation of U2-1 when variant U2-2 was absent. This up-regulation of the canonical U2-1 maintains the total U2 snRNA level in the absence of the abundant U2-2 variant (**Figures 4D** and **4E**). Northern blot analysis of all 5 major spliceosomal snRNAs confirmed constant U2 snRNA levels in wild-type and knock out cell lines (indicated by a constant ratio of U2 to U1), with no detectable change to the overall level of any snRNA, consistent with compensation of U2-2 knock-out by the up-regulation of U2-1 (**Figures 4F** and **4G**).

**Figure 4.**
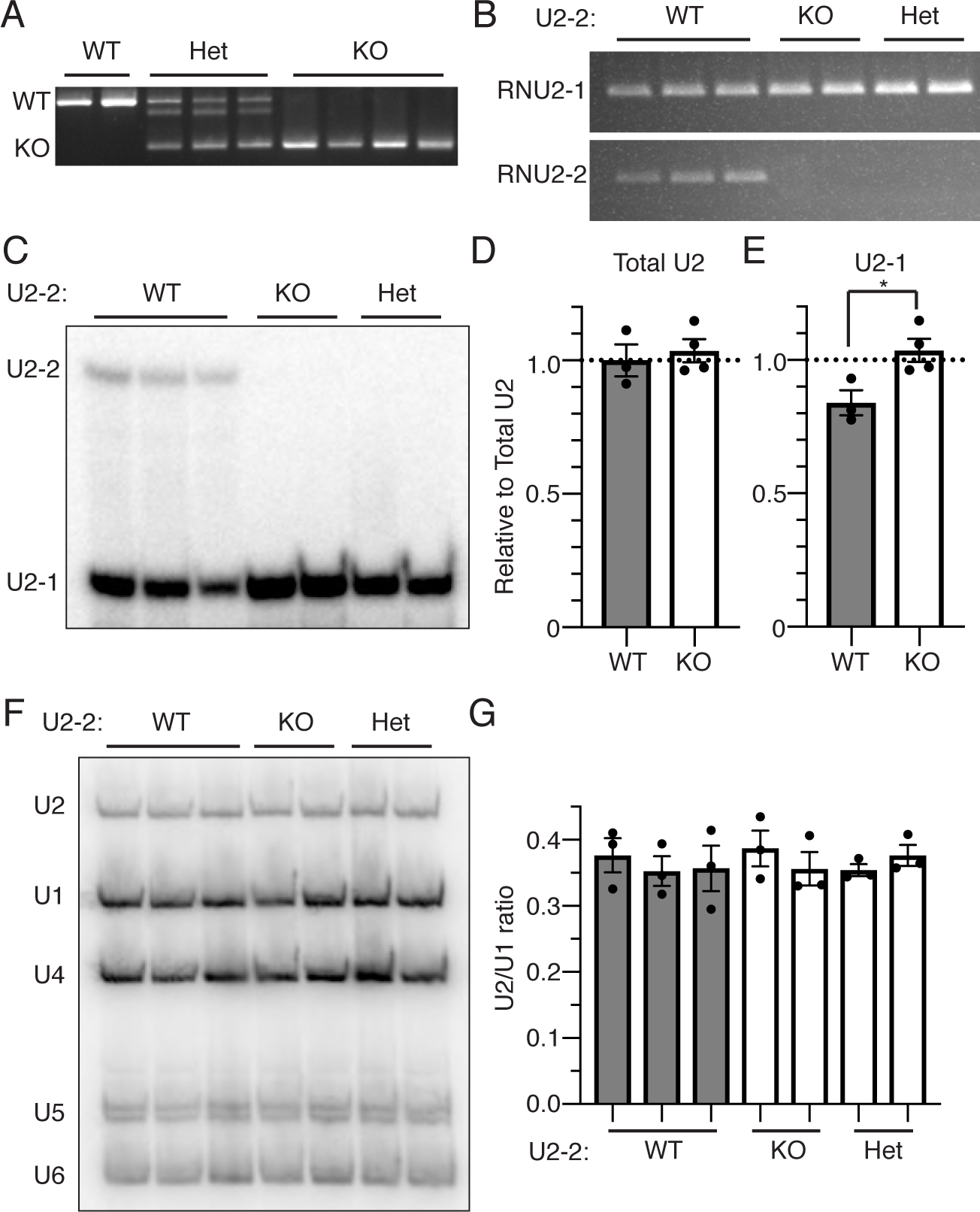
Compensatory regulation of U2-1 and U2-2 snRNAs. A) Genotyping of CRISPR/Cas9 MCF7 cell lines for deletion of U2-2 genomic loci. B) RT-PCR confirms U2-2 snRNA is not detectable in the functionally knocked out cell lines. C) Variant-specific primer extension of MCF7 U2-2^-/-^ cells suggest upregulation of U2-1 upon U2-2 knockout. D) Relative amounts of U2-1 and U2-2 detected by primer extension. Average of two technical replicates, normalized to primer intensity, were averaged for each cell line. Cell lines were averaged by genotype and normalized to average total U2 levels in U2-2^+/+^ cell lines. Error bars represent SEM of cell lines grouped by genotype. E) Quantification of canonical U2-1 in each cell line as described in (D). F) Northern blot of all snRNAs from MCF7 WT and U2-2^-/-^ suggest U2 snRNA levels are unchanged in U2-2^-/-^ cell lines, which is quantified in (G) and reported as the ratio of U2 over U1. Error bars represent SEM from 3 biological replicates. See also **Figure S4**.

### Splicing changes not detected in absence of U2-2

To determine if U2-2 knockout causes specific changes in splicing, RNA-sequencing was performed on MCF7 U2-2^+/+^ and MCF7 U2-2^-/-^ cells generated by CRISPR/Cas9 knockout (**Figure 4**) and subsequently rescued with U2-2 or U2-1 under the endogenous U2-2 promoter (**Figure 5**). In order to determine if splicing changes were U2-2-dependent, rMATS analysis of MCF7 U2-2^+/+^ and U2-2^-/-^ cells and control MCF7 cell lines from the rescue experiments described below was performed but identified few reproducible U2-2-dependent splicing changes (**Figure S5**) (Shen et al., 2014). This minimal effect on splicing caused by U2-2 loss might be expected since U2-2 represents only about 20% of total U2 snRNA expression in these cell lines and U2-2 loss is compensated for by upregulation of U2-1. Furthermore, the nucleotide differences between U2-1 and U2-2, 4 consecutive nucleotides between the Sm site and Stem loop III and a single nucleotide in stem loop IV, are in regions not known to interact with the pre-mRNA substrate or U6 snRNA during spliceosomal activation and catalysis.

**Figure 5.**
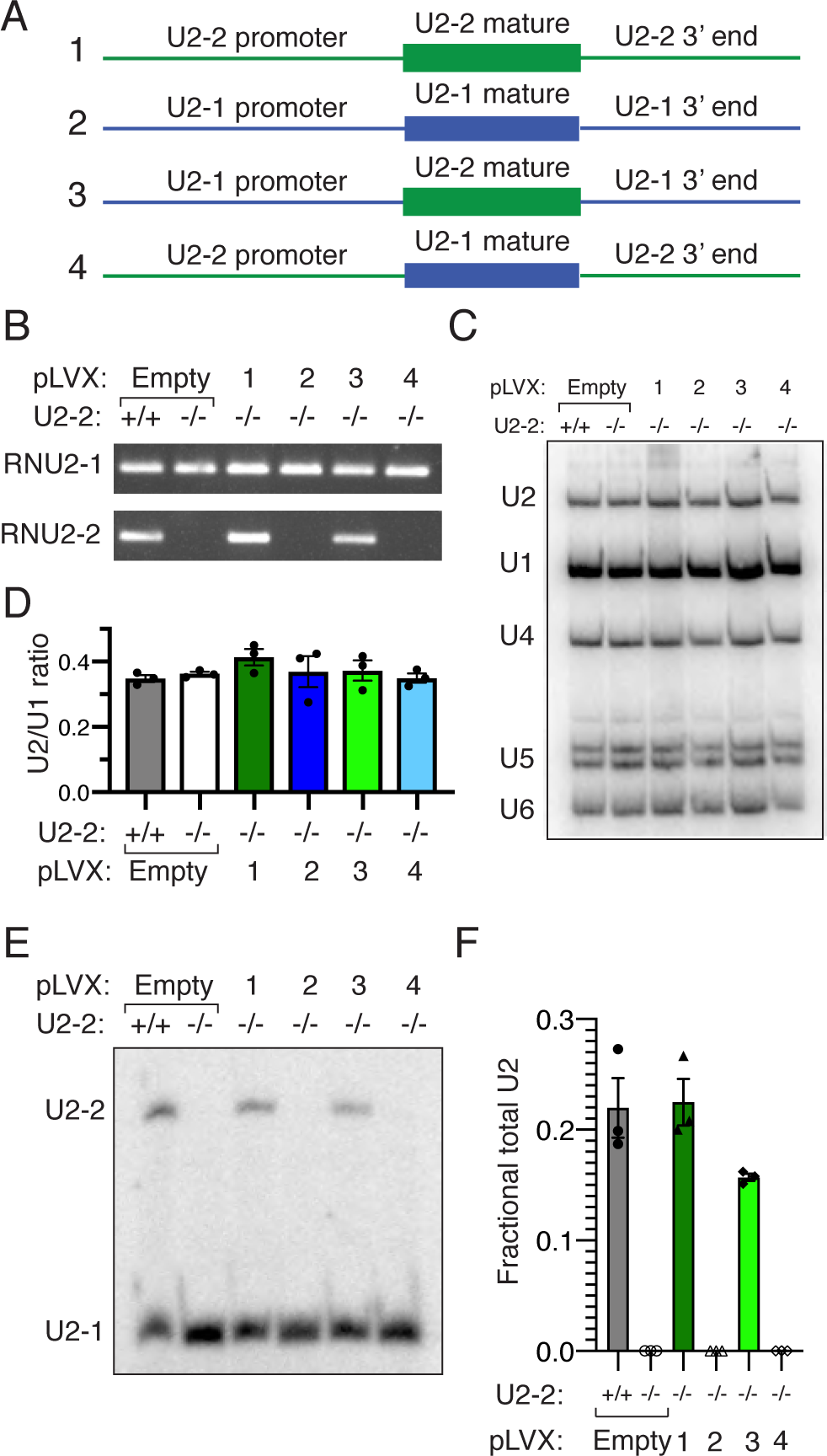
Interchangeability of U2-1 and U2-2 promoters. A) pLVX constructs designed to rescue U2-2^-/-^ MCF7 cell lines. B) RT-PCR confirmation of U2-2 snRNA expression in cell lines transfected with rescue constructs using variant specific RT-qPCR primers from Figure 2G. C) Northern blot of MCF7 U2-2 rescue cell lines. D) Quantification of total U2 snRNA level as in Figure 3F. Error bars represent SEM. E) Variant-specific primer extension of MCF7 U2-2 rescue cell lines. F) Quantification of U2-2 levels in MCF7 U2-2 rescue cell lines by RT-qPCR. Error bars represent SEM. See also **Figure S5**.

### Interchangeability of U2-1 and U2-2 promoters

To test the strength of the canonical U2-1 promoter and compare it to the variant U2-2, we made chimeric U2-2 snRNA expression vectors (**Figure 5A**). In addition to cloning the endogenous U2-1 and U2-2 genes, chimeric expression vectors where the canonical U2-1 promoter and downstream sequence drove expression of variant U2-2 snRNA and U2-1 expression driven from variant U2-2 sequences were constructed. MCF7 parental and U2-2^-/-^ cell lines were transduced with endogenous or chimeric U2 snRNA expression constructs or empty vector, and U2-2 expression was confirmed by RT-PCR (**Figure 5B**). U2-2 expression was rescued in the MCF7 U2-2^-/-^ cells when U2-2 expression was driven by either its own promoter or the U2-1 promoter.

Northern blot analysis confirmed that U2 snRNAs in all samples were processed to the proper mature length, and quantification of U2 and U1 levels detected by northern blot show that there was no significant difference in the ratio of total U2 to U1 in any of the transduced cell lines (**Figure 5C** and **5D**). We also determined the ratio of U2-2 to U2-1 by RT-qPCR and by primer extension. By RT-qPCR, U2-2 expression was rescued to wild-type levels when driven by the endogenous U2-2 promoter, and to approximately 70% of the wild-type empty vector control when driven by the U2-1 promoter (**Figure 5E**). When tested by primer extension, both wild-type empty vector and U2-2 driven by U2-1 promoter had levels greater than those detected by RT-qPCR but not statistically different from each other (**Figure 5F**). This suggests that the strength of the promoters are equal in analogous sequence contexts and that total level of U2 snRNA is regulated post-transcriptionally.

## DISCUSSION

Early identification of snRNA variants and pseudogenes was dependent on the ability to identify genomic regions with similar sequences or to separate RNA transcripts by electrophoresis. Although advancement of sequencing technologies has revealed genomic snRNA diversity in many eukaryotes, no genome-wide analysis of snRNA expression has been reported. Our bioinformatic analysis has identified transcribed snRNAs from repetitive genomic elements and discovered novel U2 snRNA variants. Levels of variant U2 snRNAs are sensitive to modulation of snRNP assembly factors suggesting snRNP assembly is a point of post-transcriptional regulation. Homeostasis upon U2-2 knockout and rescue of expression by both the U2-1 and U2-2 promoters further support that the ratio of U2-2 to U2-1 is regulated not only by transcription but also post-transcriptionally.

### Bioinformatic identification of variant snRNA transcription

Since regions flanking snRNA genes have more variation than the mature snRNA sequences, analysis of ChIP-seq data in those regions improves the probability of uniquely aligning fragments when compared to RNA-seq. Furthermore, aligning and comparing ChIP-seq datasets for histone modifications, transcription factors, and RNA pol II in multiple cell lines provides strong evidence of snRNA transcription from specific genomic loci. By allowing reads to align to multiple genomic loci, they are able to align to repetitive sequences within the snRNA genes and promoters to give a more complete picture of the transcriptional potential of each locus. However, allowing multiple alignments also increases the ChIP-seq signal of genes containing sequences identical to expressed snRNA genes. This may artifactually cause these genes to become significantly enriched. This problem would be exacerbated if only considering RNA-seq reads that align to mature sequences. In our analysis, we used hierarchical clustering and comparison to alignments that include only reads that align to unique genomic loci. This further supports our classification of snRNA genes into two groups, one transcribed and one artifactually enriched due to sequence similarity to expressed genes. While most reads are removed from unique alignments, those that remain are likely due to bioinformatic or sequencing error. However, hierarchical clustering may group genes that look truly transcribed with lower enrichment with genes enriched due to repetitive sequences. This can be observed when clustering all snRNAs genes in MCF7 cell lines (**Figure 1G**) and in genes with supposed cell-type specific transcription patterns but look to be expressed in all cell lines (**Figure S2F**).

### Potential to effect splicing and gene expression

We hypothesize that variant U2 snRNAs have the potential to affect alternative splicing through multiple mechanisms. U2 snRNA binding to the pre-mRNA substrate is essential to early 3’SS recognition. Nucleotide variation within these sequences may increase the affinity of the U2 snRNA for specific BS sequences and effect the efficiency of splicing catalysis. Additionally, U2 snRNA undergoes dramatic conformational changes as the spliceosome transitions to a more catalytically active state, and changes to sequences and/or secondary structures may alter the ability of the spliceosome to transition between different complexes, as has been shown in yeast for U2 BSL, U2 Stem loop IIa/IIb, and U6 ISL to affect splicing of sub-optimal introns (Eysmont et al., 2019; Hilliker et al., 2007; Perriman and Ares, 2007). Alternatively, spliceosomes containing U2 variant snRNPs may preferentially interact with specific metazoan-specific splicing factors to influence splice site usage.

Since the sequences in U2 snRNA known to interact with the pre-mRNA and with U6 snRNA to form the spliceosome catalytic core are identical in both U2-1 and U2-2, one might not expect intron-specific splicing changes. However, if not for the observed compensatory increase in U2-1, an overall decrease in total U2 levels could cause splicing changes similar to those previously observed upon RNase H targeted degradation by an antisense probe that targeted both U2-1 and U2-2 snRNAs (Dvinge et al., 2019). Although rMATS identified significant splicing changes between MCF7 U2-2^+/+^ and U2-2^-/-^ cell lines, few of these changes were identified in a second round of sequencing, suggesting most were due to batch or clonal effects. Also, since U2-2 levels are significantly lower in MCF7 cells than observed in many human tissues, it is possible in a different cell line or under different cellular conditions U2-2 dependent splicing changes could have been observed, possibly by altering the ability of U2 snRNP to transition between spliceosomal complexes.

### Regulation of U2 snRNA levels

Strong evidence supporting the transcription of variant snRNA genes itself does not confirm these variants participate in pre-mRNA splicing. Since U2 snRNAs are essential for multiple steps of splicing, from early recognition of the BS sequence to forming the catalytic core by providing the scaffold that presents the pre-mRNA substrates for both catalytic reactions, we further investigated the regulation and function of U2 variants. The maintenance of secondary structure by either conservative or compensatory base changes in the highly divergent U2-63 and U2-RFAM30 variants suggest an evolutionary pressure to maintain these structures. However, in the pooled human tissue RNA samples and the tested MCF7 cell line expression of these variants was low when compared to the canonical U2-1 and similar U2-2 variant. Although this might represent a significant number of transcripts per cell, it is also possible this expression level is underestimated if these U2 snRNAs are expressed more abundantly in specific cell types within these tissues.

The compensatory increase in canonical U2-1 upon U2-2 knockout might have been predicted when considering that introducing human U1 to mouse cells did not increase overall U1 levels (Mangin et al., 1985), and in differentiating cells canonical and variant U1 snRNAs are inversely expressed (Vazquez-Arango et al., 2016). It has also been observed that variant U1 sequences are able to bind multiple components of U1 snRNP but are out-competed by the canonical U1 snRNA (Somarelli et al., 2014). In human tissues where the ratio of U2-2 to U2-1 varies, U2-1 expression might be modulated by promoter accessibility or the abundancy of transcription factors. If U2-1 expression is low compared to the relative levels of assembly factors, more U2-2 is assembled into snRNPs and stabilized. This model also explains the compensatory increase in U2-1 observed in the MCF7 U2-2^-/-^ cells. Therefore, variant snRNA levels can be regulated post-transcriptionally, possibly at multiple stages of snRNP assembly or quality control where the canonical and variant snRNAs are assembled with different efficiencies.

The complex regulation of total and variant snRNA levels may be further complicated by the observed copy number variation of U2-1 in humans from approximately 5 to 82 copies (Tessereau et al., 2013; Tessereau et al., 2014). It is currently unknown how many of these repeats are transcriptionally active, how they are regulated, or if individuals with more copies express more U2-1 at the expense of U2-2. In mice, the canonical U2 snRNA is present in a multi-copy cluster, but does not have copy number variation as seen in humans. A mutation in one of five U2 snRNA genes in the cluster causes ataxia and neurodegeneration, which suggests that spatial and temporal expression of at least one specific U2 snRNA gene cannot be compensated for by the other copies (Jia et al., 2012). It remains unclear if a similar effect is possible in humans. In a mouse model of spinal muscular atrophy caused by decreased levels of SMN, a snRNP assembly factor, snRNA levels change in a tissue-specific manner (Zhang et al., 2008). Human iPS cells from spinal muscular atrophy (SMA) patients displayed dysregulation in the ratio of variant U1 to canonical U1 snRNAs (Vazquez-Arango et al., 2016). Our model suggests these findings could be due tissue-specific expression of variant snRNAs and their sensitivity to modulation in snRNP assembly factors. Lastly, mutations in multiple U2 snRNP components have been associated with various cancers and mutations to other snRNAs have been observed in various diseases and developmental disorders (reviewed in Padgett, 2012). SNPs in the canonical U1 snRNA genes are associated with various cancers (Shuai et al., 2019; Suzuki et al., 2019). Therefore, it is important to consider that disease-associated SNPs in either the canonical U2 snRNA array or expressed snRNA variants may exist but remain undetected.

## STAR METHODS

Detailed methods are provided and include the following:

- KEY RESOURCES TABLE
- LEAD CONTACT AND MATERIALS AVAILABILITY
- EXPERIMENTAL MODEL AND SUBJECT DETAILS
- METHOD DETAILS

### ○ ChIP-seq analysis

▪ Raw Fastq files were downloaded from ENCODE (https://www.encodeproject.org/) for 6 cell lines (A549,K562,HEPG2,MCF7,H1-HESC,HELA) for Pol II transcription factor and for H3K4Me3, H3K27Ac histone marks. Quality control of raw sequencing reads were performed using FastQC (*version 0.11.4*) and low quality reads were removed using cutadapt (https://cutadapt.readthedocs.io/en/stable/*; Version 1.8.3; parameters = cutadapt - q 5,10 -m 20 -o TRIMMED.fq.gz Input_fastq.gz*). The resulting quality trimmed reads from both INPUT and ChIP libraries were then aligned to custom snRNA genome using STAR aligner (*Version 2.4.2a; parameters = --runThreadN 12,-- outSAMattributes All, --outFilterMultimapNmax 20, --outFilterMismatchNmax 0, -- alignIntronMax 1*) to generate alignment (*bam*) files. deepTools (*Version 3.1.0*) with bamCompare utility (*bamCompare -b1 chip.bam -b2 input.bam -o out.bigwig -p 8 --operation log2 --binSize 25*) was used to compare ChIP with its corresponding INPUT while being simultaneously normalized for sequencing depth. For every cell line, log2(ChIP/INPUT) bigwig file is generated. Read counts for 1865 annotated snRNA genes from Gencode version 32 and RFAM version 14.1 were extracted from bam files (ChIP and INPUT) using bedtools (*Version 2.28; parameters = intersectBed with -c option*) and binomTest from edgeR package (*binomTest(y1, y2, n1=sum(y1), n2=sum(y2), p=n1/(n1+n2))*) was used to calculate the p-values for differential abundance for each snRNA between 2 libraries (INPUT and ChIP) for every cell line. Position of read was determined by adding half of the reported DNA fragment size to the 5’ end of each aligned read. snRNAs with p values <= 0.00001 and log2FC(ChIP/INPUT) > 1 for Pol II ChIP are considered to be transcribed. An R package called pheatmap was used to make clustered heatmaps for the expressed snRNA’s.

### ○ Cell culture and generation of MCF7 U2-2^-/-^ single-cell cloning

▪ MCF7 cells were a gift of X.-D. Fu (UCSD). Cells were cultured in DMEM supplemented with 10% FBS and 1x NEAA. MCF7 cells were transiently transfected with two pSpCas9(BB)-2A-GFP (PX458) vectors containing sgRNAs targeting sequences upstream and downstream of the mature U2-2 snRNA using Roche X-tremeGENE™ 9 DNA Transfection Reagent (XTG9-RO). pSpCas9(BB)-2A-GFP (PX458) was a gift from Feng Zhang (Addgene plasmid #48138; http://n2t.net/addgene:48138; RRID:Addgene_48138). After 48 hours, transfected cells were single-cell sorted into 96-well plates using BD FACSaria II. DNA from clonal cells was isolated using Zymo Quick DNA miniprep w/ Zymo-spin IIC columns or Epicenter QuickExtract DNA Extraction Solution (QE09050). PCR primers flanking the sgRNA targets were using for genotyping utilizing NEB Taq 2x Master Mix (M0207). sgRNA and primer sequences are listed in Supplementary **Table S4**.

### ○ Plasmids

▪ The CMV promoter was excised from pLVX-puro lentiviral expression plasmid (Clonetech). pLVX-puro was digested with ClaI and XhoI, blunted with NEB Quick Blunting kit (E1201S), and ligated with NEB T4 DNA ligase (M0202S) to generate pLVX-dCMV. Endogenous U2-1 and U2-2 genes were amplified from human genomic DNA. The U2-2 promoter and 5’ end of the mature sequence were amplified separately from the 3’ end of the mature and downstream sequence. Overlapping PCR of these fragments substituted GUGUAGUA with an AscI restriction site (GGCGCGCC). The endogenous U2-1 and U2-2 genes with AscI substitution were cloned into Invitrogen Zero Blunt Topo vector (pCR4Blunt-TOPO vector - K287540). The pCR4Blunt-TOPO plasmids containing both U2-1 and U2-2 sequences and pLVX-dCMV were digested with EcoRI. The inserts containing the U2 snRNA sequences were then ligated into the linearized pLVX-dCMV and confirmed by Sanger sequencing. The U2-2 expression construct with AscI substitution was then digested with AscI, and an NEB HiFi assembly kit was used to replace the AscI site with endogenous GUGUAGUA sequence. Chimeric constructs, flanked by ClaI and EcoRI restrictions sites, were synthesized by GeneWiz and were cloned into pLVX. PCR primer sequences for amplification of U2-1 and U2-2 are listed in Supplementary **Table S4**.

### ○ Generation of MCF7 U2-2^-/-^ rescue cell lines

▪ Rescue MCF7 U2-2^-/-^ cell lines were generated using the pLVX-puro lentiviral system with the plasmids generated as described above. Lentiviral infections were performed using X-tremeGENE™ 9 DNA Transfection Reagent (Roche). 48 hours post infection, the cells were selected with 2 µg/ml puromycin.

### ○ RNA isolation and quantification

▪ RNA from cell lines grown to 90% confluence was isolated using Trizol according to manufacturer’s protocol. RNA was quantified first on Nanodrop2000 and diluted to either 500 ng/ul or 250 ng/ul. Final concentration of RNA was then determined on a Denovix QFX fluorometer using Qubit RNA BR Assay kit according to the manufacturer’s protocol.

### ○ RT-qPCR

▪ RNA was isolated as described above. cDNA from 500 ng of total RNA was reverse transcribed using Invitrogen Superscript III reverse transcriptase with 50 pmol of random hexamer primers in a 20 ul reaction, according to manufacturer’s protocol. Relative snRNA levels were determined by RT-qPCR. snRNA qPCR primers and diluted cDNA were added to Applied Biosystems™ Power™ SYBR™ Green Master Mix for 10 ul reaction in 384 well plate. qPCR was performed on Applied Biosystems ViiA 7 Real Time PCR system. After 10 minutes of incubation at 95°C, 40 cycles of amplification (95 °C for 15 s, 60 °C for 1 minute) were measured and melting point analysis was performed to ensure specificity. The efficiency-corrected threshold cycle (ΔCT) method was used to determine the relative levels of snRNA. Primer sequences are listed in Supplementary **Table S4**.

### ○ Primer extension

▪ Antisense DNA oligo (Supplementary **Table S4**) hybridizing to both U2-1 and U2-2 was 5’-end labeled with gamma ^32^P-ATP using Invitrogen T4 PNK and gel purified on 20% PAGE-urea gel. A thin slice of gel was crushed in a 1.5 ml Eppendorf tube with 250 ul of H2O, 25 ul 3M NaAc, and 300ul saturated phenol (pH 4.3 ± 0.2). The slice was vortexed at 37°C for one hour and centrifuged 15,000xg for 15 mins. The aqueous layer was collected and phenol chloroform extracted, and precipitated using two volumes ethanol overnight at -20°C. The solution was centrifuged at 15,000 g for 15 mins at 4°C, the pellet washed with 75% ethanol, air-dried, and resuspended in H2O.

▪ 1.2 ul of 5x annealing buffer was added to 250,000 CPM of resuspended gel-purified 5’-end labeled DNA oligo and 250 ng of total RNA to a final volume of 6 ul in 1x annealing buffer (final concentration: 50mM Tris pH 8.3, 10 mM DTT, 60 mM NaCl). The annealing reaction was heated to 90°C for 3 mins in a heat block before being removed, placed on the bench at room temperature and allowed to cool to 55°C. The annealing reactions were immediately placed in an ice slurry. 9 ul of the RT reaction mixture [0.5 ul 5x RT buffer (250 mM Tris pH8.3, 50 mM DTT, 300 mM NaCl, 150 mM Mg acetate), 3ul 1x AB buffer, 5x dNTP Mix (2.5 mM ddGTP, dATP, dCTP, dTTP in 1x AB), 0.5 ul of Superscript III RT diluted 1:25 in Dilution Buffer (200 mM KPO4 pH 7.2, 2 mM DTT, 0.2% Triton X-100, 50% glyerol)] was then added to the 6 ul annealed RNA. The RT reaction was incubated for 5 minutes in a 37°C water bath followed by 30 minutes in a 42°C water bath. The RT reaction was then precipitated overnight at -20°C in 1 ml of 100% ethanol after the addition of 20 ug of glycogen carrier and 2 ul of 3 M NaOAc pH 5.5. The precipitated reaction was then centrifuged at 15,000xg, washed with 75% ethanol, and excess supernatant was pipetted off the pellet. The pellet was resuspended in 6 ul RNA loading buffer (6. urea, 1xTBE, 10% glycerol) and loaded onto a 15% PAGE-urea gel. The gel was transferred to GE BAS-IP MS 2040 E phosphor-imaging screen and imaging was obtained on Fuji Typhoon 9400 imaging system.

### ○ Northern blotting

▪ Total RNA was isolated from cell lines at 90% confluence as described above. 1 ug total RNA was added to 8 ul RNA loading buffer and loaded onto an 8% PAGE-urea gel. RNA was transferred from the gel to GE Hybond+ nitrocellulose membrane in 0.5xTBE and crosslinked using a Stratalinker UV crosslinker (120 mJoules). Antisense oligos for U1, U2, U4, U5, and U6 were 5’ radiolabeled with Invitrogen T4 PNK and added to hybridization buffer (0.75 M NaCl, 0.1 M NaHPO4, 7% SDS, 1x Denhardt’s solution and 100 ng/ml sonicated salmon sperm DNA), hybridized to membrane overnight and washed x2 with wash buffer (40 mM NaHPO4, 2% SDS, 1 mM EDTA). Membrane was transferred to GE BAS-IP MS 2040 E phosphor-imaging screen and image was obtained on Fuji Typhoon 9400 imaging system. Primer sequences are listed in Supplementary **Table S4**.

### ○ RNA-seq and bioinformatic analysis of alternative splicing

▪ Total RNA was isolated from 3 consecutive passages of MCF7 parental cell lines and from 3 clonal MCF7 U2-2^-/-^ cell lines. Total RNA was also isolated from 3 consecutive passages MCF7 parental cells and MCF7 U2-2^-/-^ both transfected with pLVX-dCMV. RNA was bioanalyzed and submitted to Novogene for sequencing. RNA was poly(A) selected and sequenced to a depth of 40 million reads per sample. Raw Fastq files were obtained from NOVOGENE and was checked for quality of reads with FastQC (version 0.11.4). The raw reads for both bulk-control and U2-2 knock downs were aligned with a splice aware aligner such as STAR (Version 2.4.2a; parameters = --runThreadN 12 --outSAMattributes All -- outSAMtype BAM SortedByCoordinate --outSAMstrandField intronMotif -- outFilterMultimapNmax 1 --outFilterMismatchNmax 6 --alignIntronMax 50000 -- sjdbGTFfile gencode.gtf --quantMode TranscriptomeSAM GeneCounts -- twopassMode Basic --outFilterScoreMinOverLread 0.41 -- outFilterMatchNminOverLread 0.41). Multivariate Analysis of Transcript Splicing with replicates (rMATS: Version 3.2.5: command = python RNASeq-MATS.py -b1 bulk1.bam,buk2.bam -b2 ko1.bam,ko2.bam -gtf gencode.v27.annotation.gtf -o rMATS_U_3.2.5_Bulk_vs_KO -t paired -len 150 -analysis U) was used to detect differential splicing events. The program outputs 5 main category of events: Skipped Exons (SE), Retained Introns (RI), MXE (Mutually Exclusive Exons), A3SS (Alternative 3’ splice site) and A5SS (Alternative 5’ splice site). Custom based Perl scripts were used to filter events which have FDR >= 0.01 and junction counts <=20 junction reads. The remaining events were called significant.

## ● QUANTIFICATION AND STATISTICAL ANALYSIS

### ○ Primer extension

▪ ImageJ was used to quantify both total and relative amounts of U2-1 and U2-2. At least two technical replicates normalized to the amount of free primer were averaged to determine U2-1 and U2-2 snRNA levels. The ratio of U2-2 to U2-1 snRNA was calculated and then averaged per genotype. The unpaired, two-tailed Student’s t test was applied. Results equal to or above a 95% confidence interval (p value < 0.05) were considered statistically significant. Unless p values are specifically identified, they are notated as follows: * = 0.01 to <0.05, ** = 0.001 to <0.01, *** = 0.0001 to <0.001. Unless otherwise specified, error bars represent standard error of the mean (SEM).

### ○ RT-qPCR

▪ Technical duplicates were and three biological replicates were performed per target and sample. The efficiency-corrected threshold cycle (ΔCT) method was used to determine levels. The fraction of total U2 for each snRNA was calculated by dividing the level of each snRNA by the sum of U2-1 and U2-2 and then averaged per sample. The unpaired, two-tailed Student’s t test was applied as described above.

## ● DATA AND CODE AVAILABILITY

○ GEO Accession numbers
○ Unprocessed images

## SUPPLEMENTAL INFORMATION

Supplemental Information consists of 5 figures and 4 tables.

## ACKNOWLEDGMENTS

We are grateful to Matt Gamble, Teresa Bowman, and Magda Konarska for helpful discussions and critical readings of the manuscript. We thank David Shechter (Einstein) and X.-D. Fu (UCSD) for generous gifts of cells lines, and members of the Flow Cytometry Core Facility for expert assistance (NCI P30CA013330). This work was supported by NIH grant GM57829 to C.C.Q and a Cancer Center Support (core) grant from the NCI to AECOM. C.C.Q is a scholar of the Irma T. Hirschl Trust.

## AUTHOR CONTRIBUTIONS

Conceptualization, B.K. and C.Q. Methodology and Investigation, B.K. Software, V.G. Data curation and Analysis, B.K. and V.G. Writing, B.K., C.Q., and V.G. Funding acquisition and Supervision, C.Q.

## DECLARATION OF INTERESTS

The authors declare no competing interests.

## SUPPLEMENTAL FIGURE LEGENDS

**Figure S1.**
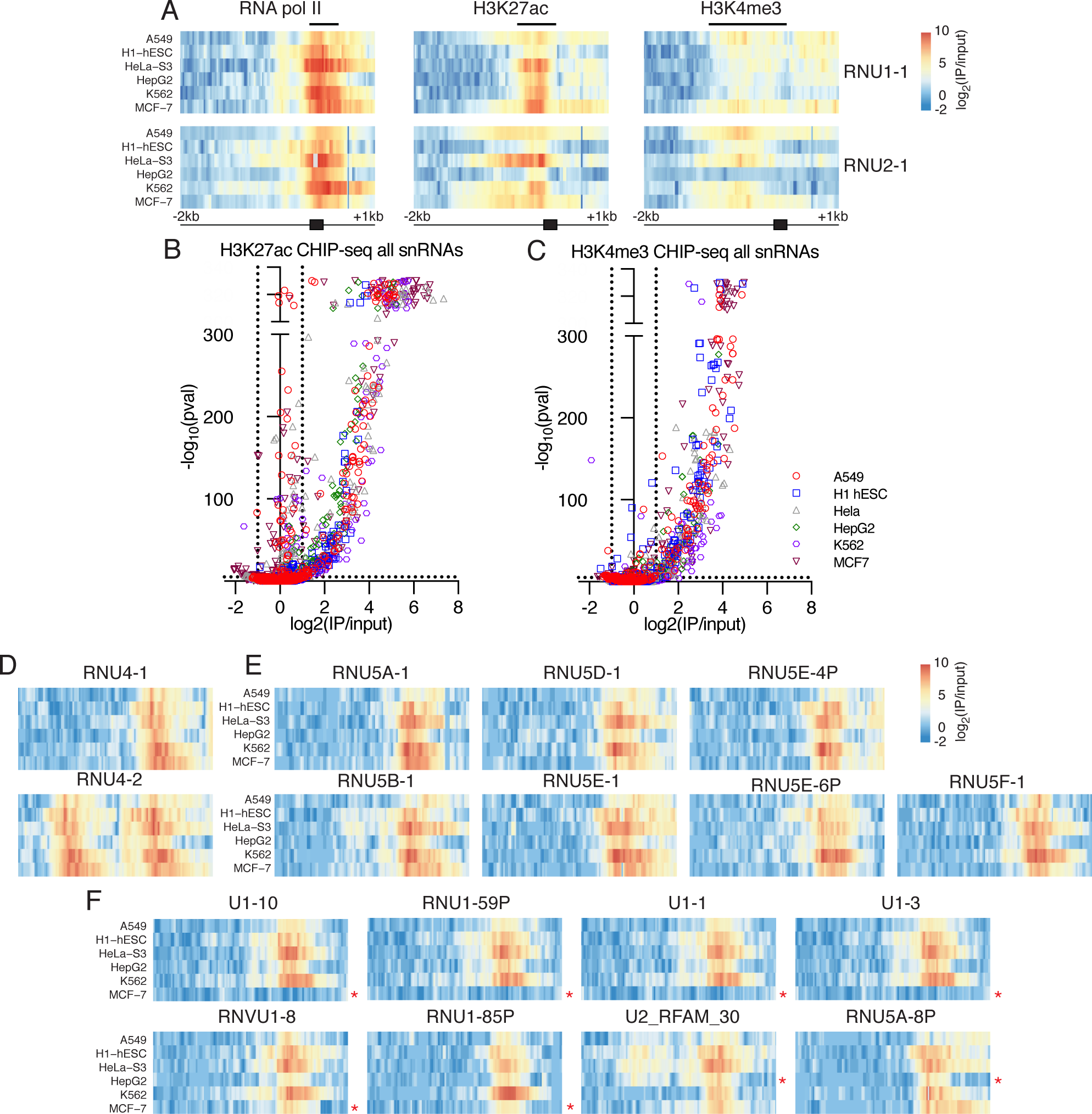
Determination of actively transcribed snRNA genes, Related to Figure 1. (A) RNA Pol II, H3K27ac, and H3H4me3 ChIP-seq analysis of canonical U1 and U2 snRNA genes, RNU1-1 and RNU2-1, reveals regions enriched for each target as indicated by black bar above the graph (Pol II from -50 bp to +400 bp, H3K27ac from -400 bp to +200 bp, and H3K4me3 from -1000 bp to +200 bp relative to the transcription start site (TSS)). Each row is either RNU1-1 or RNU2-1 as indicated on right from a different cell line as labeled on left. Each row includes 2 kb upstream of the TSS and 1 kb downstream. The TSS and approximate length of a mature snRNA (200 bp) are indicated by the black box. (B) Volcano plot shows significant enrichment of H3K27ac density over specific snRNA genes in six common cell lines. log2(IP/input) on x-axis shows magnitude of enrichment while -log10(p-val) is calculated from binomial test and plotted on y-axis. All -log10(p-val) greater than 300 are shown above break in graph. Jitter (noise) added to values >300 for visualization. Cut-offs of >1 for log2(IP/input) and >5 for -log10(p-value) for significance are indicated by dotted lines. (C) Volcano plot shows significant enrichment of H3K4me3 density over specific snRNA genes in six common cell lines. log2(IP/input) on x-axis shows magnitude of enrichment while -log10(p-val) is calculated from binomial test and plotted on y-axis. All -log10(p-val) greater than 300 are shown above break in graph. Jitter (noise) added to values >300 for visualization. Cut-offs of >1 for log2(IP/input) and >5 for -log10(p-value) for significance are indicated by dotted lines. (D) ChIP-seq Pol II heatmaps of U4 and (E). U5 snRNA genes identified as transcribed in Figure 1D across six cell lines as indicated in labels to right of (D) U4 and left of (E) U5 snRNA. Each row is as described in **Figure S1A**. (F) snRNA variants that are differentially grouped in six tested cell lines by hierarchical clustering reveals four U1 variant snRNA genes absent from MCF7 cells (top row) and those that still may be transcribed despite hierarchical clustering. Red (*) indicates cell lines where variant does not cluster with other expressed snRNA genes.

**Figure S2.**
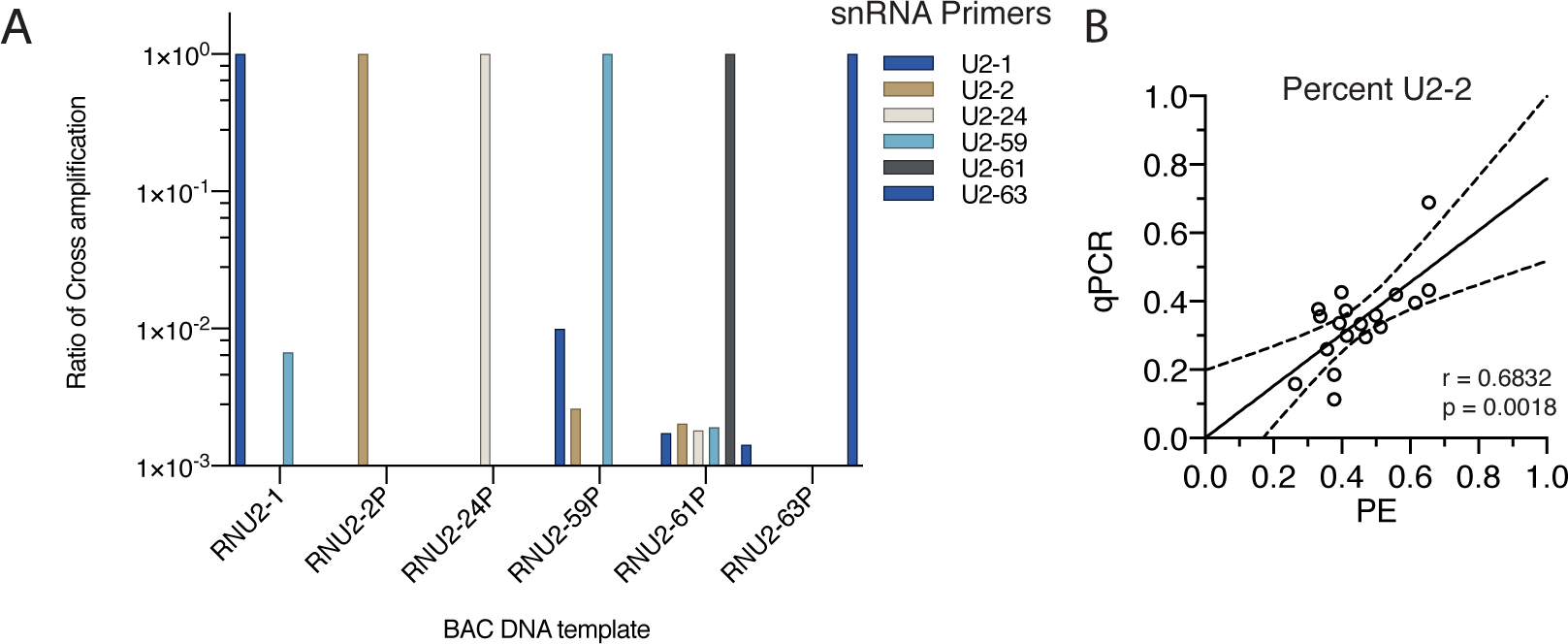
Specific detection of U2 snRNA variants, Related to Figure 2. (A) RT-qPCR primer cross reactivity. RT-qPCR on BAC DNA templates grouped on x-axis shows ability to amplify with specific primers, each a different color. Scale in log10. (B) Correlation between ratio of U2-2 to total U2 snRNA levels as detected by RT-qPCR and variant-specific primer extension. r and p values calculated using Pearson correlation. 95% confidence interval shown by dotted lines. Although there is a significant correlation between the primer extension assay and RT-qPCR, primer extension utilizes a single primer and only requires a single reaction; in contrast, RT-qPCR may introduce and amplify any bias caused by the sequence variation (U2-1 and U2-2 qPCR reverse primers contain the sequence variation at their 3’ ends), resulting here in an underestimation of U2-2 levels. Thus, we consider variant-specific primer extension to be the more reliable assay.

**Figure S3.**
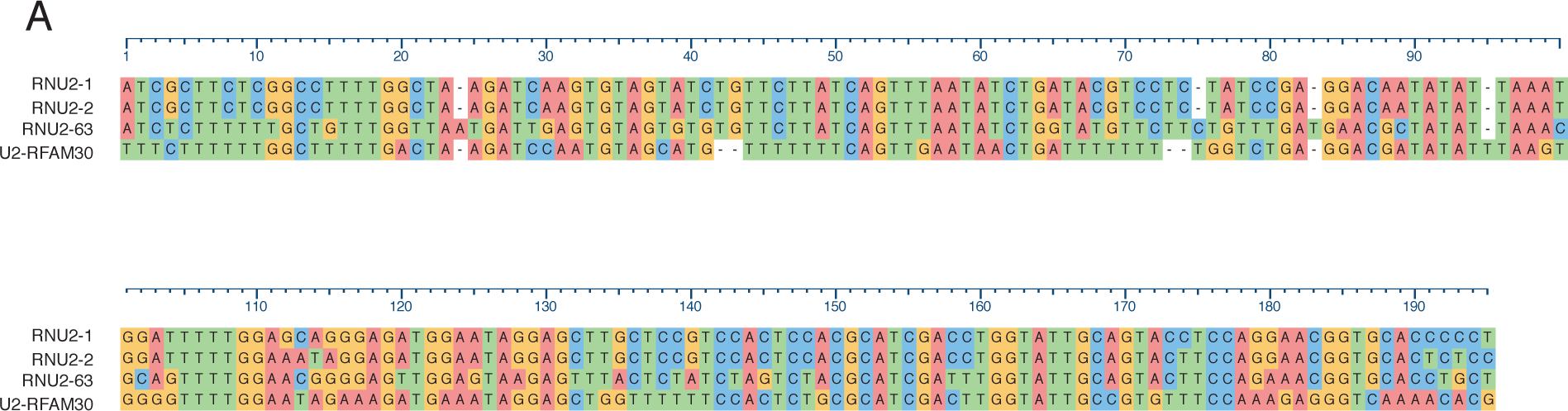
Clustal Omega alignment of transcribed U2 snRNAs, Related to Figure 3. (A) Clustal Omega alignment (Sievers et al., 2011) of four annotated U2 snRNA with enriched RNA pol II density.

**Figure S4.**
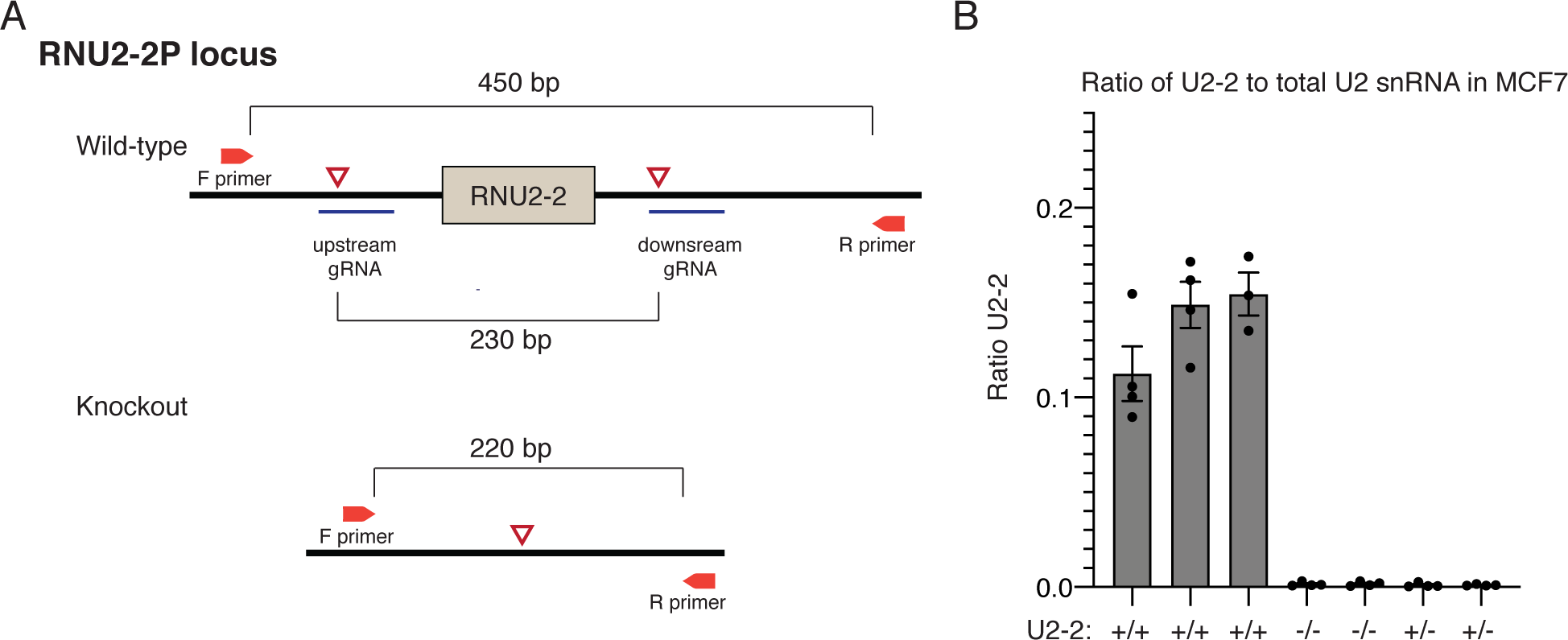
CRISPR/Cas9 knock out of U2-2, Related to Figure 4. (A) Diagram of CRISPR gRNAs targeting either side of the mature RNU2-2 sequence and positioning of genotyping primers. (B) Ratio of U2-2 to total U2 snRNA levels by RT-qPCR in MCF7 wild-type and knockout cell lines. Error bars represent SEM of three biological replicates.

**Figure S5.**
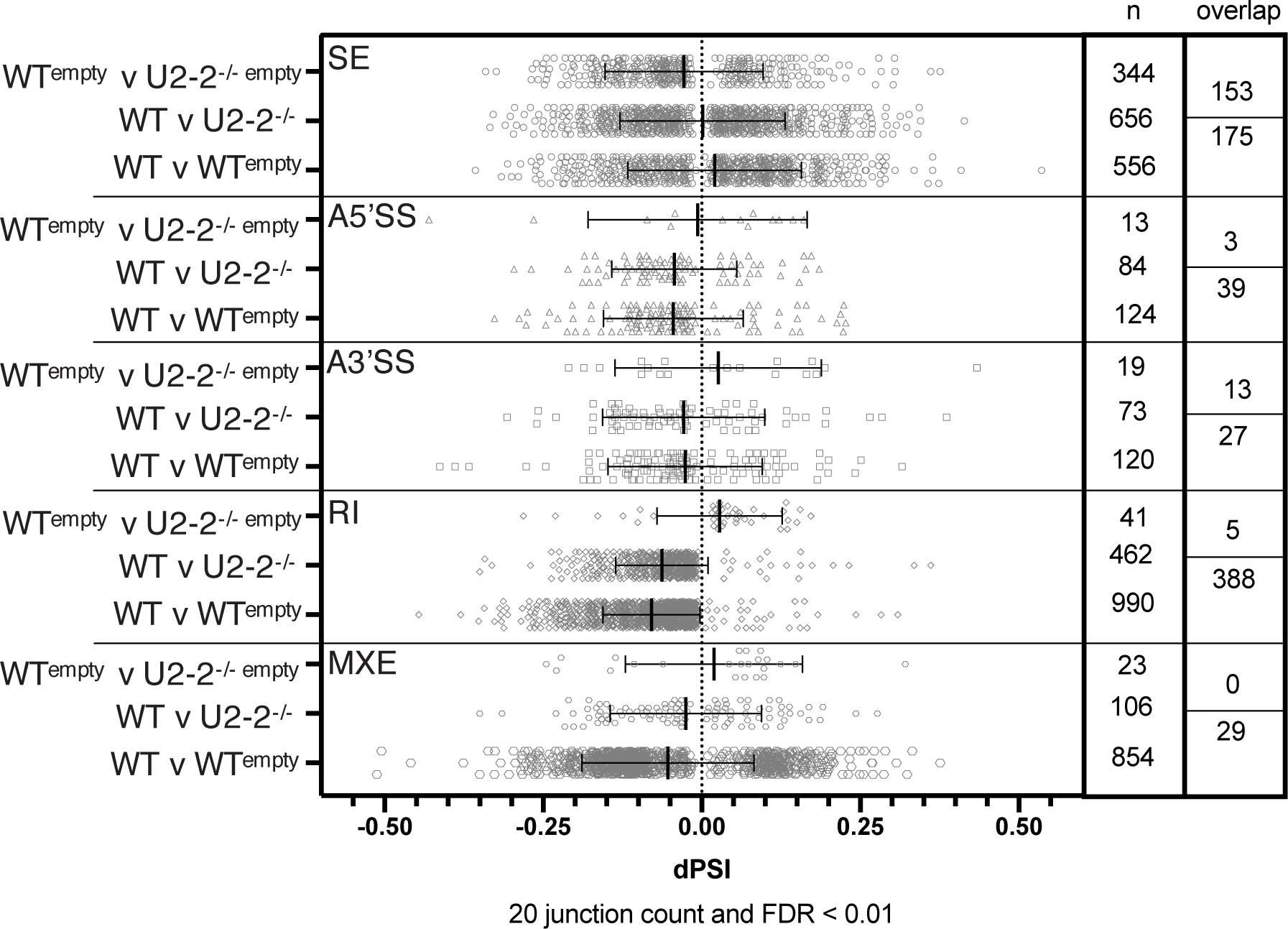
Nominal splicing changes in MCF7 U2-2^-/-^ cells, Related to Figure 5. Significant splicing changes identified in rMATS analysis of poly(A) selected RNA-seq. Number of total significant alternative splicing events and overlap of those events between two replicates of MCF7 U2-2^+/+^ and MCF7 U2-2^-/-^ shown on columns to the right. More overlap in events between two MCF7 U2-2^+/+^ replicates than between MCF7 U2-2^+/+^ and MCF7 U2-2^-/-^ suggest identified differences are not dependent on U2-2 expression.

**Supplemental Table S1.**

| Species | Genome Size (MB) | Alternative Splicing | Annotated snRNA loci | U1 snRNA | U2 snRNA | U4 snRNA | U5 snRNA | U6 snRNA |
| --- | --- | --- | --- | --- | --- | --- | --- | --- |
| S. cerevisiae | 12.1 | <1% | 5 | 1 | 1 | 1 | 1 | 1 |
| D. melanogaster | 165 | 50% | 47 | 14 | 7 | 4 | 8 | 5 |
| C. elegans | 100 | 31% | 79 | 13 | 22 | 6 | 13 | 25 |
| D. rerio | 1400 | 46% | 1160 | 371 | 18 | 30 | 145 | 586 |
| M. musculus | 3300 | 78% | 1296 | 198 | 128 | 44 | 13 | 872 |
| H. sapiens | 3300 | 85% | 1966 | 176 | 172 | 93 | 34 | 1414 |

